# Expression profiling of single cells and patient cohorts identifies multiple immunosuppressive pathways and an altered NK cell phenotype in glioblastoma

**DOI:** 10.1101/792846

**Authors:** Helen J. Close, Lucy F. Stead, Jérémie Nsengimana, Katrina A. Reilly, Alastair Droop, Heiko Wurdak, Ryan K. Mathew, Robert Corns, Julia Newton-Bishop, Alan A. Melcher, Susan C. Short, Graham P. Cook, Erica B. Wilson

## Abstract

Glioblastoma (GBM) is an aggressive cancer with a very poor prognosis. Generally viewed as weakly immunogenic, GBM responds poorly to current immunotherapies. To better understand this problem we used a combination of NK cell functional assays together with gene and protein expression profiling to define the NK cell response to GBM and explore immunosuppression in the GBM microenvironment. In addition, we used transcriptome data from patient cohorts to classify GBM according to immunological profiles. We show that glioma stem-like cells, a source of post-treatment tumour recurrence, express multiple immunomodulatory cell surface molecules and are targeted in preference to normal neural progenitor cells by natural killer (NK) cells *ex vivo*. In contrast, GBM-infiltrating NK cells express reduced levels of activation receptors within the tumour microenvironment, with hallmarks of TGF-β mediated inhibition. This NK cell inhibition is accompanied by expression of mutiple immune checkpoint molecules on T cells. Single cell transcriptomics demonstrated that both tumour and haematopoietic-derived cells in GBM express multiple, diverse mediators of immune evasion. Despite this, immunome analysis across a patient cohort identifies a spectrum of immunological activity in GBM, with active immunity marked by co-expression of immune effector molecules and feedback inhibitory mechanisms. Our data show that GBM is recognised by the immune system but that anti-tumour immunity is restrained by multiple immunosuppressive pathways, some of which operate in the healthy brain. The presence of immune activity in a subset of patients suggests that these patients will more likely benefit from combination immunotherapies directed against multiple immunosuppressive pathways.

## Introduction

Glioblastoma (GBM) is the most common and aggressive type of primary adult brain cancer. Current treatments include debulking neurosurgery and adjuvant chemo/radiotherapy. Despite these therapies, median overall survival is just 12-24 months (1). Recent developments in cancer immunotherapy provide one potential approach to improve patient outcomes (2). However, despite significant therapeutic impact on several solid tumour types, immune checkpoint blockade (ICB) is yet to demonstrate benefit in GBM treatment (3).

Evasion of host immunity is a hallmark of cancer (4). Tumours exploit the negative feedback mechanisms that the healthy immune system uses to dampen immune responses. These mechanisms include the recruitment of immune cells with suppressive activity, the expression of immunosuppressive cytokines, such as Transforming Growth Factor (TGF)-β, and immune checkpoints including Programmed cell Death (PD)-1 and Cytotoxic T Lymphocyte Antigen (CTLA)-4 (5). Chronic interactions between tumour cells and infiltating T cells leads to an exhausted phenotype, an unresponsive but reversible state with an altered transcriptional profile (6). Exhausted tumour infiltrating lymphocytes express immune checkpoints, and antibodies that target these molecules can reinvigorate anti-tumour immunity (7).

Mutations in the tumour genome are a source of neoantigens and mutation frequency is a surrogate marker for immunogenicity (8). For several tumours, neoantigen load is correlated with survival and with response to immune checkpoint blockade; melanoma and lung adenocarcinoma have higher mutational load, greater T cell infiltration, greater PD-1 expression and consequently show better responses to anti-PD-1 therapy (9). Compared to other solid tumours, mutation frequency and T cell infiltration levels in GBM are low. However, GBM-infiltrating T cells have been isolated against a number of germline encoded antigens overexpressed in the tumour, indicating that T cell responses are at least possible (10). Glioblastoma cells are a target for natural killer (NK) cells, however the number of GBM-infiltrating NK cells is also low (11). The paucity of NK cells and T cells in GBM is compounded by the high proportion of suppressive myeloid-lineage cells (12) able suppress to lymphocyte function (5).

Importantly, GBM-infiltrating T cells express PD-1 (13), yet reports of initial anti-PD-1 (Nivolumab) clinical trials are not encouraging (2). This indicates that PD-1 expression *per se* is not sufficient to allow responsiveness to therapy and that additional suppressive components of the GBM immune landscape regulate many effectors of anti-tumour immunity.

Here we show that *in vitro*, GBM cells are recognised and killed by NK cells, however NK cells derived from GBM tumours have an altered cell surface phenotype consistent with their inhibition in the tumour microenvironment. We have explored the basis of this inhibition and identify numerous immunosupressive mechanisms operating in GBM contributed by both tumour and immune cell compartments. These immunosuppressive pathways, some of which appear to operate in the normal brain, are a barrier to effective immunotherapy, but also represent candidate therapeutic targets to reinvigorate tumour NK cell interactions.

## Materials and Methods

### Ethics statement

Ethical approval for this study was granted by the Ethics committee at the Leeds Teaching Hospitals NHS Trust, Leeds UK (REC number 10-H1306-7).

### Classification of GBM patients Consensus Immune Cluster (CIC)

CIC classification (14) was applied to GBM tumour transcriptome data from TCGA. Briefly, consensus cluster analysis of melanomas, used the expression of 380 genes specific to 24 immune cell types ((15), this produced 6 subtypes which we termed CICs. The average expression of each gene within each CIC is the cluster centroid. Using TCGA data, we used the nearest centroid method (16) to classify each GBM tumour into one of the CICs according to the highest Spearman correlation coefficient with the centroids. For each of the 24 immune cell types of the immunome compendium (Bindea et al. 2013), we calculated a score per GBM tumour, graphically represented using a heatmap.

### Differential expression of genes in GBM CIC2 and CIC4 and in REMBRANDT data

To compare expression of selected genes in different patient groups we used RNAseq data from TCGA (obtained via TCIA (7)), assigning patients to either CIC2 or CIC4 (as above). In addition, we used microarray data from the REMBRANDT study (17) downloaded from Betastasis.com. For REMBRANDT, patient samples were classified as GZMA high expressors or GZMA low expressors based on the median expression value (n=214). Expression of selected genes was compared between CIC2 and CIC4 (for TCGA data) or the GZMA^high^ and GZMA^low^ patients (for REMBRANDT) and analysed using non-parametric, unpaired statistical testing (using GraphPad Prism).

### Single cell data and normal brain analysis

Single cell (sc)RNAseq data was downloaded from (18) and expression of candidate genes analysed. Data was visualised using rStudio Version 1.0.143 (package: gplots 3.0.1) using heatmap.2. Untransformed data clustering (unsupervised) was performed (Euclidean distance). Individual cells were classified as “tumour” or “immune” according to co-expression of SOX9 and EGFR(18) and PTPRC respectively. For all genes expression of >0 was scored positive. For immune (PTPRC+) and tumour (SOX9+EGFR+) cells, the number of different immunomodulatory molecules expressed was counted for each cell and the percentage of immune and non-immune cells expressing immunomodulatory genes plotted. Expression of individual genes in non-tumour bearing brain tissue was downloaded from (19). This data is also available (with graphical output) at BrainRNAseq.org.

### Tumour tissue and blood, collection and processing

After ethical approval and informed consent, tumours were resected and stored in PBS, or within the cavitron ultrasonic surgical aspirator (CUSA) (20) Samples were washed in PBS, CUSA samples were prepared as (20), all samples were filtered through 40 µm cell strainer, washed twice PBS, centrifuged 400 × g, 5 mins and resuspended in PBS, 0.5% BSA, 0.05% sodium azide. Matched patient blood was diluted with PBS, layered over Ficoll (Axis-Shield PoC, Oslo, Norway) and centrifuged at 800 × g for 20 mins. Tumour and blood derived cell were stained with appropriate antibodies (see supplementary methods table).

### Primary cells and cell lines

Neural progenitor cells (NP1) were isolated from a patient undergoing surgery to treat epilepsy (21). The primary lines, GBM1 and NP1 were generated at the Scripps Institute. GBM11, GBM13 and GBM20 were derived at the University of Leeds using the same method and culture conditions (22). PBMC were isolated from whole blood of healthy donors as above. NK cells further separated using an NK cell isolation kit (Miltenyi Biotec, Germany), and cultured in DMEM supplemented with 10% FBS, 10% human AB serum (Sigma-Aldrich).

### Surface antigen screening

GSC cell lines were harvested using 0.25% trypsin/EDTA and fluorescently labelled for 60 mins at 37°C and 5% CO_2_, in serum-free media with one of the following cell dyes: 0.4µM cell tracker™ (CT)- green CMFDA (488nm excitation), 2µM CTorange-CMRA (488nm excitation), 2µM CTviolet-BMQC (407nm excitation) or 5µM calcein blue-AM (407nm excitation) (all from Invitrogen.) All populations were washed 3 times, mixed together, and plated at a density of 1×10^6^ total cells/well in 96-well round bottom plates (Nunc). Cells were stained as per the manufacturer’s instructions with 242 antibodies from the BD Bioscience Lyoplate screening panel, followed by Zombie NIR (Biolegend) for 30 mins before resuspension and analysis by flow cytometry. Cells were gated based on their emitting fluorescence at 520nm (CTgreen loaded), 580nm (CTorange loaded), 540nm (CTviolet loaded) or 449nm (calcien blue loaded). The median fluorescence intensity (MFI) for each gated population, for each antigen and isotype control emission at 668nm (Alexa647 emission) was generated and GSC lines scored as positive if more than 20% of the population expressed the antigen. Flow cytometric analysis was performed using FacsDiva (BD), FlowJo (Treestar) and Kaluza (Beckman Coulter) software.

### Natural killer cytotoxicity assays

Target tumour cell lines were labelled with the relevant cell dye (see surface screen) for 1hr at 37°C, washed twice and plated at 2×10^5^/well. NK cells were pre-activated with 20ng/ml IL-15 for 48hrs and mixed with targets at the E:T ratios indicated. After 5 hours, cells were pelleted (300 × g, 5 mins), washed with PBS and stained with Zombie NIR (Biolegend), for 15 mins at room temperature. Competitive cytotoxicity assays were set up as above, the two target cell types under test (GBM and neural progenitors) were labelled with either CTgreen or CTviolet, mixed 1:1 and used as a target population at an E:T of 5:1.

## Results

### Glioma stem-like cells are effective targets of NK cells

Effective therapy for GBM will require the elimination of the radioresistant GBM stem-like cells (GSCs) that are largely responsible for recurrence (23). Whilst tumour associated antigen-specific T cells offer a highly selective therapeutic approach, antigen-independent effector cells, such as NK cells, have the potential to target and destroy GBM tumour cells that have a low neoantigen load.

We used three patient derived GSC lines (described in Polson et al., 2018) shown to exhibit a stem cell-like expression profile and recapitulate high-grade gliomas in orthopic xenograft mouse models (22, 24), and performed cytotoxicity assays, using peripheral blood derived, IL-15 activated NK cells to confirm NK cell mediated killing. Tumour cells differentiated from GSCs are more sensitive to NK cell killing than the GSC themselves (26) but GSCs are killed by NK cells in the presence of activating cytokines (Figure 1A) (25). We further tested whether NK cells activated with IL-15 would be efficient killers of GSCs but retain specificity for GSCs over normal neural progenitor cells. We performed an NK cytotoxicity assay using a mixed target cell population comprised of tumour GSC cells and normal neural progenitor (NP) cells (22) at a ratio of 1:1. For all donors, IL-15 activated NK cells killed tumour cells in preference to the NP cells (Figure 1B). These results suggest that, in short term *in vitro* cultures when sufficient immune cells are present and activated, GSCs are an effective and preferential target for NK cells.

**Figure 1.**
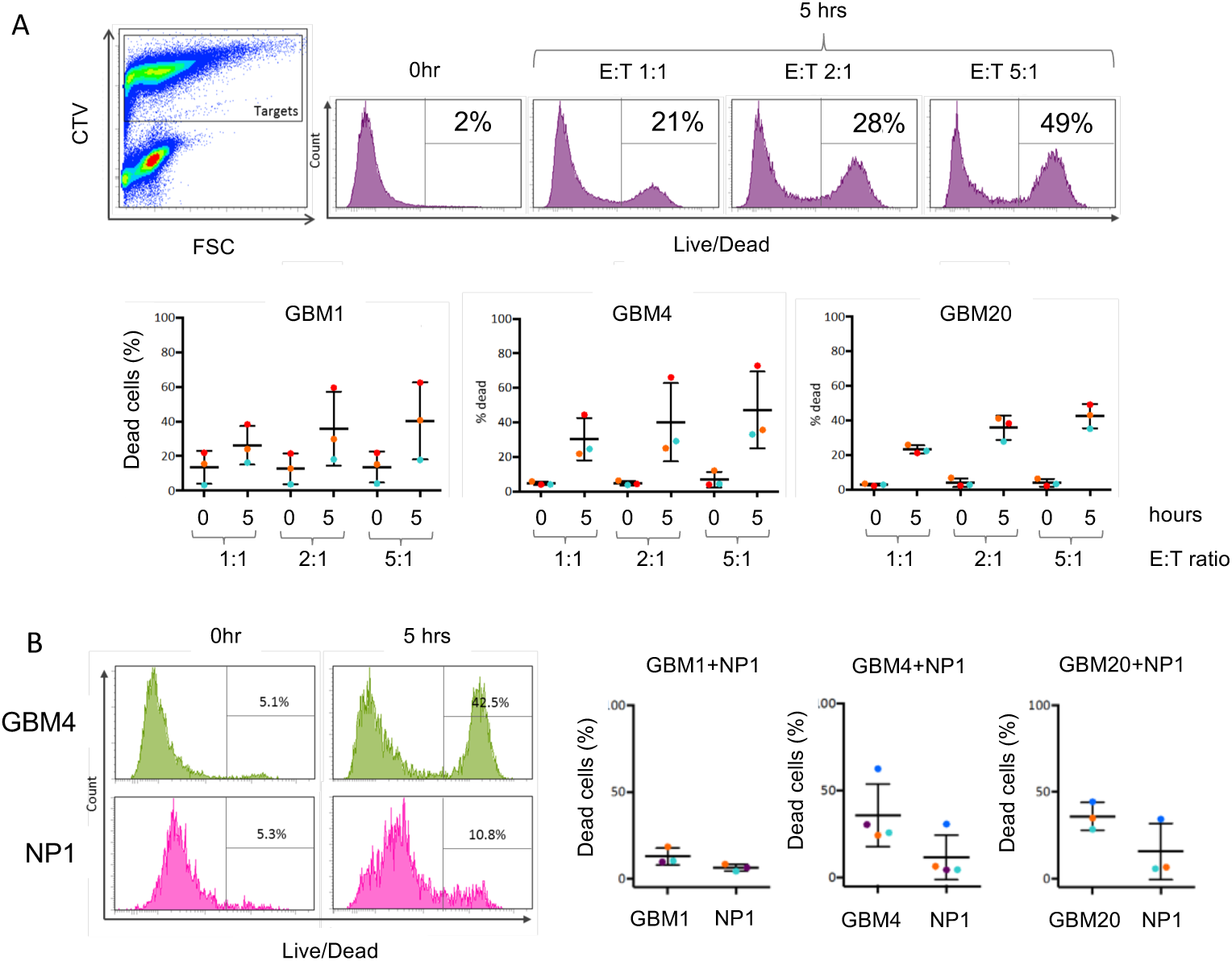
NK cell-mediated killing of glioma stem-like cells. **A)** NK cell cytotoxicity: Cell tracker violet-labelled GSC lines (targets) were co-cultured with unlabelled, IL-15 activated NK cells (effectors) for five hours at effector:target (E:T) ratios as shown. Co-cultures were then stained with a live/dead discriminator. The panel on the left shows identification of effector and target cells in the co-culture (for gating purposes) and the panels to the right show death of the labelled target cells at the different E:T ratios. The zero hour control is included as background cell death of the GSC cells. The three graphs summarise data obtained using three GSC lines (GBM1, GBM4 and GBM20) and three different NK cell donors (coloured dots), with standard deviation from mean. **B)** NK cell specificity: Cytotoxicity assays of IL-15 activated NK cells co-cultured with a 1:1 mix of the GSC line (indicated) and neural progenitor cells (NP). The GSC and NP lines were labelled with different cell tracker dyes allowing their fate in the assay to be determined separately. The flow cytometry plots show the percentage of dead GSC (here GBM4) and NP cells after zero and five hours of co-culture with NK cells. The graphs summarise this data for assays containing the three GSC lines using NK cells from four separate donors (coloured dots), with standard deviation from mean.

### Patient-derived NK cells exhibit an altered cell surface phenotype in GBM

The presence of infiltrating NK cells in GBM (11) coupled with their ability to recognise and kill GSCs (Figure 1) suggests that they are rendered non-functional in the GBM tumour microenvironment. We performed flow cytometry-based analysis of intratumoural NK from GBM tissue and compared their surface phenotype to NK cells derived from autologous peripheral blood as well as blood from healthy donors. NK cell populations were defined as NKp46^+^ and CD3^-^ due to high expression of CD56 (NCAM-1) on GBM tumour cells within the sample (Supplementary Figure 1). To confirm sampling of immune cells from within GBM tumour tissue (and not from blood contamination of the tumour sample) we assayed the expression of PD-1 on T cells, and showed significantly enhanced expression of PD1 on tumour-derived T cells compared to their blood counterparts (Figure 2A). Expression levels of NK cell surface molecules were similar on the blood-derived NK cells from both healthy donors and GBM patients. However, the expression of the tumour-sensing NK cell activating receptors NKp30, NKG2D and DNAM-1, and the surface molecules tetherin/CD317 and CD2 were all significantly reduced on the GBM tumour-derived NK cells compared to those from matched peripheral blood (Figure 2B). Together with higher expression of PD-1 on GBM derived T cells compared to matched peripheral blood (Figure 2A), we also found higher expression of LAG-3 and CTLA-4 (although differences in CTLA-4 expression did not reach statistical significance) (Supplementary Figure 2).

**Figure 2.**
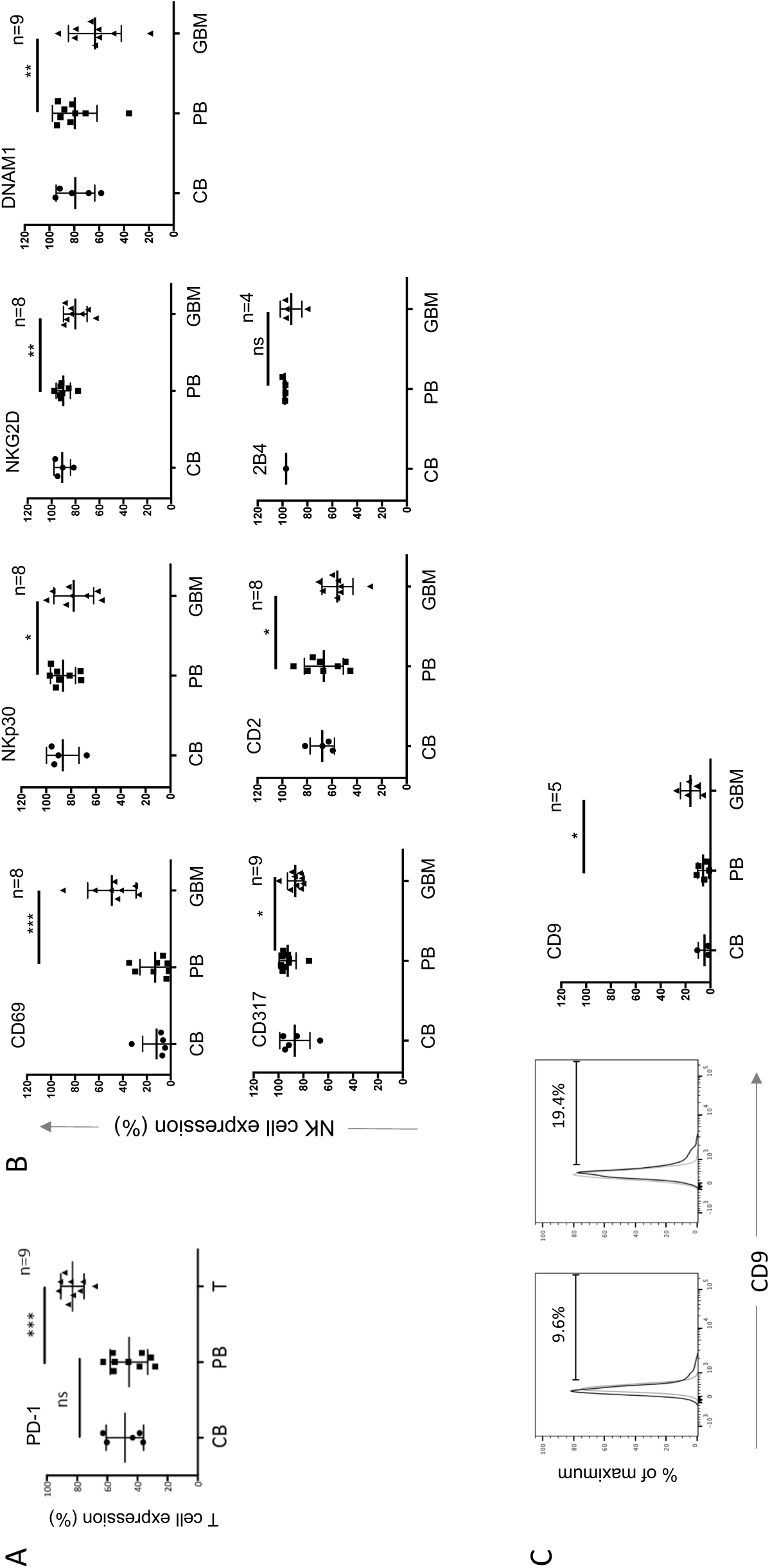
The cell surface phenotype of GBM-infiltrating lymphocytes. **A)** Expression of PD-1 on CD3+ T cells in GBM patient tumour (GBM), patient blood (PB) and control blood from healthy donors (CB). Each dot represents a single patient sample (n is the number of GBM patient samples analysed); the bar indicates the mean± standard deviation. The patient-derived tumour (GBM) and blood (PB) samples were analysed using a paired t test; *P<0.05, **P<0.01; ***P<0.001; ns: not significant. **B)** Expression of NK cell surface molecules (gating on CD45+, NKp46+, CD3^neg^ cells) in GBM patient tumour, patient blood and control blood from healthy donors as in (A). **C)** Representative histograms of CD9 expression on PB and GBM derived NK cells, grouped data as in (A).

Members of the TGF-β family are highly expressed in GBM and are important in maintaining the GSC pool (25). Furthermore, we and others have previously shown that TGF-β reduces the expression of NKp30, NKG2D and DNAM-1 on NK cells and is associated with their functional inactivation (26, 27). Importantly, TGF-β induces the expression of the tetraspanin CD9 on NK cells (28) and we detected significantly increased expression of CD9 on the surface of the GBM-resident NK cells compared to NK cells from matched peripheral blood (Figure 2C). The reduced expression of NK cell activating receptors coupled with the increased expression of CD9 is suggestive of TGF-β mediated evasion of NK cell cytotoxicity in the GBM microenvironment.

Collectively, these data demonstrate that GBM resident immune effector cells are constrained by at least two separate mechanisms, the reduced expression of NK cell activation receptors and the increased expression of immune checkpoint molecules on T cells.

### Surface antigen screening of GSCs identifies candidate immunomodulatory molecules

The GSC lines are selectively targeted by NK cells *in vitro* but evade NK cells and other immune effector cells *in vivo*. To understand what immunomodulatory molecules expressed by GSC might be responsible for immune activation and inhibition we analysed GSCs for the expression of cell surface immunomodulatory molecules. Using a flow cytometry-based screen, we identified 116 cell surface antigens expressed on four GSC lines lines (Supplementary Table 1). Molecules detected on the GSCs included those associated with the cancer stem cell phenotype (CD24, CD44 and CD90) (Figure 3A), as well as widely expressed cell surface molecules, such as MHC class I (and β2-microglobulin), CD71 and CD98, as expected. Several immune inhibitory molecules were highly expressed, such as the immune checkpoint ligands PD-L1 (CD274) and PD-L2 (CD273), providing a source for inhibition of PD-1 expressing T cells (Figure 3A). In addition, we found expression of the ectonucleotidase CD73, that together with CD39 generates extracellular adenosine to inhibit both NK cells and T cells via purinergic receptors (29), as well as expression of CD200 and CD47, modulators of myeloid cell activity (Figure 3A). Ligands of NK cell activation receptors, such as MICA/B (NKG2D ligand) and CD112 (a DNAM-1 ligand), as well as CD80 (a T cell costimulator) were detected, along with CD54 (ICAM-1) and CD50 (ICAM-3); ligands of LFA-1 required for NK cell and T cell mediated cytotoxicity (30). The GSC cell surface screen therefore revealed expression of a repertoire of targetable cell surface molecules with the potential to activate and inhibit NK cells, T cells and myeloid cells. This prompted us to explore the expression of immunosuppressive pathways in more detail, using a publicly available GBM single cell gene expression dataset (18). Amongst 3589 single cells, we identified 757 co-expressing SOX2 and EGFR (defined by Darmanis *et al* as tumour cells) and 1527 cells expressing PTPRC (encoding CD45, a marker of cells of haematopoietic origin.) We next performed unsupervised hierarchial clustering using expression of lineage marker genes and genes encoding candidate immunosuppressive functions, which identified two main groups; non-immune (comprising tumour and neuronal cells) and immune cells (PTPRC+) (Figure 3B). The immune cell group was dominated by expression of numerous myeloid cell markers (Figure 3B). Genes encoding immunosuppressive functions were expressed within both the immune and non-immune clusters (Figure 3B) and overall, individual immune cells expressed a greater number of immunosuppressive genes than tumour cells (Supplementary Figure 3A). Consistent with the altered cell surface phenotype of GBM-resident NK cells (Figure 2A), we found widespread expression of TGFB family transcripts accounted for by TGFB1 expression in the myeloid cells and TGFB2 and TGFB3 expression in non-immune cells. Furthermore, HLAG (which plays a key role in regulating NK cell activity in pregnancy and cancer (31, 32)) was also widely expressed. The HLA-G protein inhibits myeloid cells via receptors LILRB1 and LILRB2 (31), both of which were expressed in the immune compartment at the mRNA level. We identified strong expression of the receptor-ligand pair HAVCR1 (TIM3) and LGALS9 in the myeloid cluster (92% of immune cells expressed HAVCR2 or LGALS9 and 60% expressed both genes; Supplementary Figure 3B). This scRNA-seq data along with the GSC surface antigen screen shows that both tumour and immune infiltrating cells express receptors and ligands that together constitute a complex network of immunosuppression. For example, the combined action of CD73 and CD39 generate immunosuppressive adenosine (29); our data shows expression of CD73 by the tumour cells (Figure 3A) and NT5E (encoding CD39) by the immune fraction (Figure 3B), with Mohme et al demonstrating CD39 expression by GBM infiltrating T cells (13).

**Figure 3.**
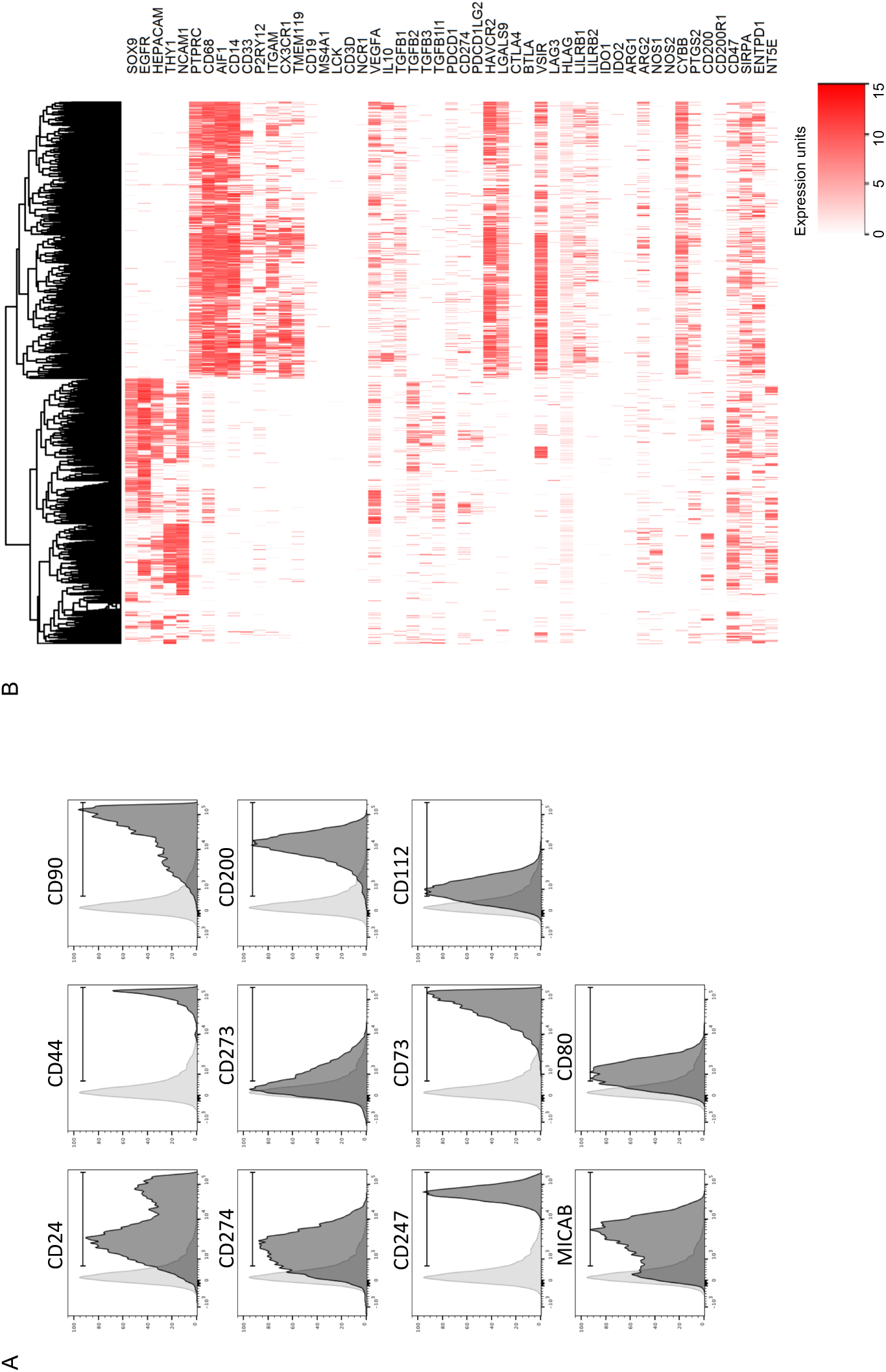
The repertoire of immunosuppressive molecules expressed in GBM. **A)** Expression of selected cell surface antigens on GSC lines, the data shows expression by GBM20. A summary of expression across the four GSC lines is provided in supplementary table 1. **B)** Single cell (sc) RNAseq data (18) was clustered, revealing immune and tumour groups marked by PTPRC and SOX9/EGFR co-expression respectively. Expression of marker genes for cell lineages and those encoding immunomodulatory molecules are indicated. Expression is scored according to the values and key shown.

Furthermore, to assess whether this immunosuppressive network was induced in response to tumour, we analysed gene expression data derived from normal brain tissue (33) (Supplementary Figure 4). Microglia/macrophage (the only cell population in the normal brain expressing PTPRC/CD45), constitutively express several immunosuppressive genes, including immune checkpoints VSIR and HAVCR2, and checkpoint ligands LGALS9, CD274 and PDCD1LG2. Some components of the immunosuppressive network found in brain tumours are therefore present in the healthy brain.

**Figure 4.**
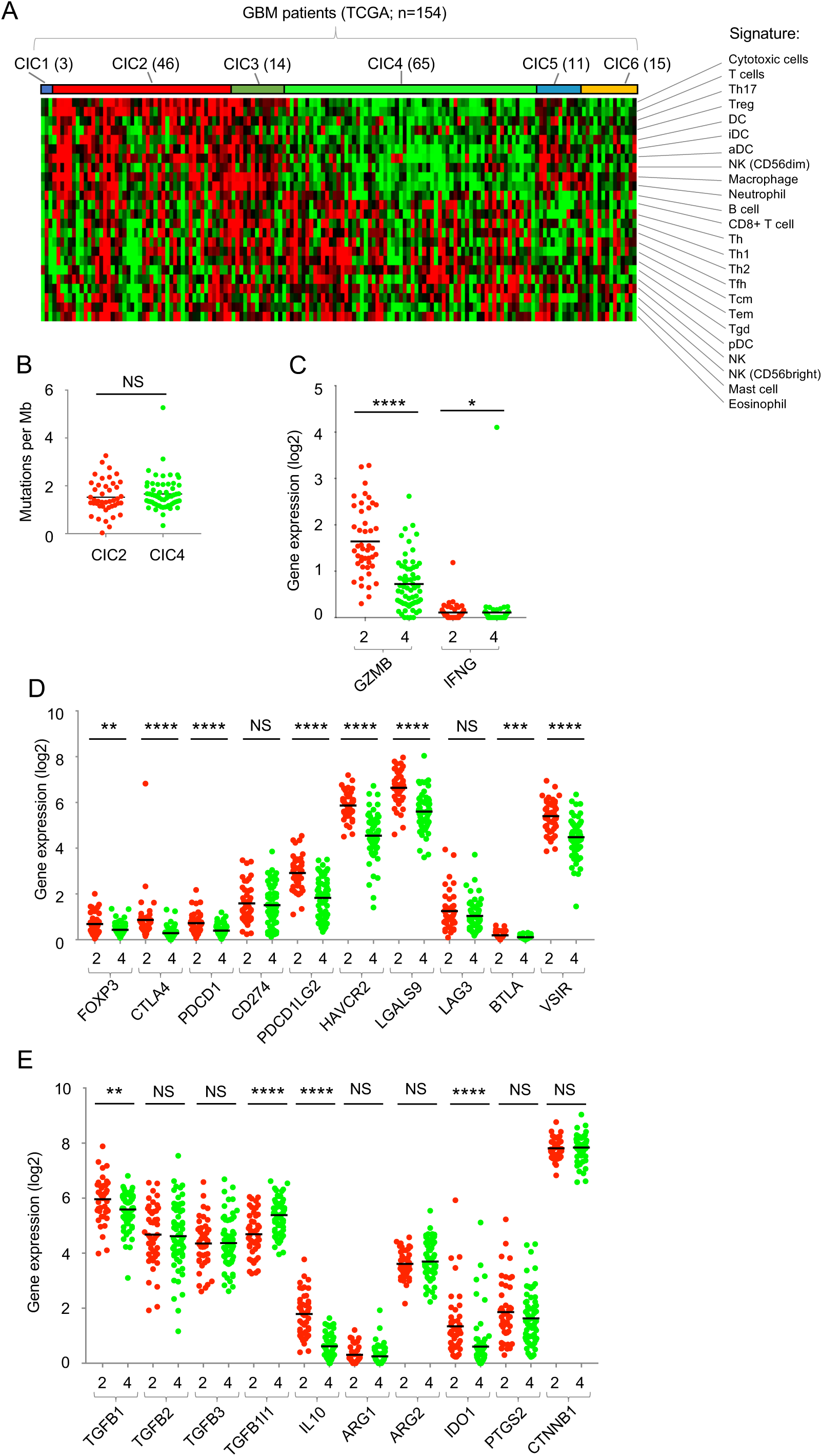
A spectrum of immune activity in GBM. **A)** Classification of GBM tumours (from 154 patients in the TCGA dataset) into consensus immunome clusters (CIC) using the nearest centroid classification. The number of patients in each CIC is indicated in brackets. The cell signatures used to derive the CIC (14) are shown. **B)** Mutational load in GBM CIC2 (red) and CIC4 (green) expressed as mutations per megabase. **C)** Expression of GZMB and IFNG in CIC2 (red) and CIC4 (green). **D)** Expression of negative regulators of immunity in CIC2 (red) and CIC4 (green). **E)** Expression of cytokines and enzymes associated with immunosuppressive activity. For panels B-E, data from CIC2 and CIC4 were compared using the Mann Whitney test; NS: not significant, *P<0.05,**P<0.01,***P<0.001,****P<0.0001.

### A spectrum of immune activity in GBM patients

We have previously used tumour transcriptome data to cluster melanoma patients according to their immune cell infiltrate (14). This approach identified six Consensus Immunome Clusters (CICs), with one cluster enriched in cytotoxic cells (CIC2), and another (CIC4) having low immune infiltrates and significantly worse survival (14). We used this approach to classify GBM transcriptome data (from TCGA) and, like the situation in melanoma, the cohort of 154 patients clustered into the six CICs (Figure 4A), with 2 main clusters CIC2 (high immune infiltrate) and CIC4 (low immune infiltrate). We found that CIC2 was significantly enriched for tumours of the mesenchymal subtype (34) (Supplementary Table 2) that has been previously shown to have prolonged survival (21). However, unlike melanoma (14), immune infiltration (reflected in the CIC clusters) was not associated with significant differences in survival in GBM (Supplementary Figure 5). There was also no significant difference in mutation burden, a surrogate of neoantigen load reflecting immunogenicity, (8, 9) between CIC2 and CIC4 in GBM (Figure 4B). These data demonstrate that patients can be stratified based on the immune infiltrate but that, unlike melanoma, this stratification has no effect on patient outcomes under the conditions of treatment currently employed.

Immune activation induces feedback inhibitory pathways, including the expression of immune checkpoint molecules, and we therefore attempted to use the expression of genes in these pathways to understand the immune environment within the GBM CIC clusters. To do this we compared expression of anti-tumour effector functions and immunomodulatory genes in CIC2 and CIC4. Expression of the GZMB and IFNG genes were significantly higher in CIC2 than CIC4, consistent with the increased infiltration of cytotoxic T cells and NK cells (Figure 4C). Furthermore, genes encoding immune checkpoint molecules (CTLA4, PDCD1, HAVCR2, BTLA and VSIR), their ligands (PDCD1LG2, LGALS9) and FOXP3 were also expressed at significantly higher levels in CIC2 than CIC4 (Figure 4D), as were genes encoding soluble mediators of immunosuppression such IL10, TGFB1 and IDO1 (Figure 4E). To confirm this we used microarray data from the REMBRANDT study (17) and GZMA gene expression as a simple surrogate for immune infiltration (35). This analysis confirmed the significantly higher expression of multiple immunosuppressive functions in patients with increased expression of anti-tumour effector functions (Supplementary Figure 6). Collectively these data drive our understanding of the GBM immune microenvironment, demonstrating a spectrum of immune infiltration, functionally compromised by an active immune-inhibitory network.

## Discussion

Our analysis demonstrates tumour and immune mediated immunosuppression within the GBM tumour microenvironment, functionally inactivating GBM anti-tumour immunity. We demonstrate reduced expression of tumour-sensing activating receptors on GBM-resident NK cells consistent with TGF-β activity (26). The TGF-β family cytokines play a manifold role in glioma progression, including maintenance of the GSC pool, proliferation, invasion, angiogenesis and immunosuppression (36). Multiple mechanisms of tumour-mediated downregulation of NK cell activation receptors have been identified. However, we favour TGF-β as a modulator of the NK cell phenotype in GBM, as we show reduced expression of activation receptors coupled with increased expression of CD9, a tetraspanin induced by TGF-β in NK cells (28). Mohme et al showed that infiltrating T cells expressed PD-1, TIM-3 and CD39 (13) characteristic of T cell exhaustion (37). Our analysis extended these findings by identifying CTLA-4 and LAG-3 on GBM-infiltrating T cells. Thus, GBM-infiltrating NK cells have reduced expression of activating receptors, whereas T cells have increased expression of immune checkpoint molecules, resulting in inhibition of both classes of cytotoxic lymphocytes. Furthermore, we identified CD73 on the GSC cell surface and together with CD39 on infiltrating T cells, these ectonucleosidases may act together to generate immunosupressive adenosine which inhibits both NK cells and T cells (29). Similar to Castriconi *et al* (38), we demonstrate that activated NK cells are capable of recognising and killing GSC cell lines *in vitro*, and we further show that NK cells discriminate between the GSC and a normal neural progenitor cell line.

The analysis of NK cells in GBM and their interaction with GSCs led to a more extensive analysis of the immunosuppressive network. Our data highlights the abundance of immunosuppressive pathways operating in GBM. The most abundant immune cells in GBM belong to the myeloid lineage (12) and we show that GSCs express cell surface molecules such as PD-L1 and CD47, which inhibit phagocytosis by macrophage (18, 39, 40). These data demonstrate that immune inhibition within GBM is mediated by both immune and non-immune lineages cooperating to provide a pro-tumour environment. Interestingly, CD47 and CD200 are important regulators of microglial activity and brain inflammation in non-malignant disease (41). Several of the immunosuppresive pathways evident in GBM are in place in the normal brain. In response to TGF-β, microglia suppress immunological activity and promote normal microglial functions such as synaptic pruning and neuronal growth support (42). The expression of genes such as HAVCR2, its ligand LGALS9 and CD274, along with TGFB2 and IL10 safeguard the normal brain against excessive inflammation (43). Muller et al demonstrate that infiltrating macrophages rather than resident microglia encode immunosuppressive cytokines within the GBM microenvironment (44). Thus, immunosuppressive pathways operating in the GBM-free brain are utilised and extended upon by infiltrating myeloid cells, contributing to the extensive immunosuppressive network.

The identification of a spectrum of immune infiltration across the GBM cohort, accompanied by evidence of anti-tumour effector function and feedback inhibitory pathways suggests that GBM should not simply be regarded as an immunogenically “cold” tumour. The high expression of mutiple feedback inhibitors along with high expression of GZMB and IFNG in CIC2 identifies ongoing, or at least prior, immune activation in a subset of GBM patients, restrained by the action of these inhibitory pathways. Indeed, melanoma shows inter-patient heterogeneity of immune responses and regarding melanoma as “hot” fails to account for this variability. In melanoma, immune heterogeneity impacts upon survival (14) and the success of immunotherapy, with high expression of PD-1, CD8+ T cell infiltration and higher mutational burden associating with response to therapy (9). We found no evidence for differential survival in GBM according to our CIC classifications, suggesting that the extensive immunosuppressive network removes any impact of immune control of GBM progression. However, this survival data is based on standard therapy and, by analogy with melanoma, we suggest that GBM CIC2 patients are more likely to respond to immunotherapy. Moreover, the extensive immunosuppressive network suggests that targeting multiple inhibitory pathways will be a likely requirement of GBM immunotherapy.

The mechanisms underlying inter-patient heterogeneity of immune response in GBM are unclear (45). β-catenin mediated immune evasion pathways operate in CIC4, the group with the poorest prognosis in melanoma (14), whereas here we found no evidence of CTNNB1 differential expression between CIC2 and 4 and neither was their mutational burden significantly different. However, GBM arises through various combinations of oncogene and tumour suppressor mutations and several of these genes regulate tumour immune responses (46, 47). Thus, differences in oncogene and/or tumour suppressor gene mutations between patients is one potential factor underlying the spectrum of immune activity seen across the GBM cohort.

GSCs are effective targets of NK cells *ex vivo*, but GBM infiltrating NK cells have a surface phenotype bearing the hallmarks of TGF-β mediated immunosuppression. Further exploration of immunosuppressive pathways using gene and protein profiling indicated that both tumour and immune cell components contribute inhibitory factors. These pathways are a barrier to effective immunotherapy, but also represent candidate therapeutic targets. Combined checkpoint blockade is already outperforming monotherapy in melanoma (46). Targeting combinations of the multiple immune checkpoints (PD-1, LAG-3, CTLA-4, TIM-3, VSIR/VISTA) or other immunosuppressive molecules (e.g. ectonucleosidases, TGF-B, IL-10) may prove beneficial in GBM. Strategies to activate NK cells *in situ*, e.g. via the use of oncolytic viruses (48-50) and methods to alter the immune composition of GBM will also benefit from alleviating key immunosuppressive pathways. Importantly, GBM is not universally devoid of immune activity and a subset of patients with evidence of immune activity suggests that combinatorial immunotherapy would be most effective when patients are stratified acoording to immune infiltrates.

## Acknowledgements

This work was supported by Cancer Research UK, The Brain Tumour Charity, Brain Tumor Research and Support across Yorkshire and the University of Leeds. We are indebted to those investigators who have made large datasets and analysis tools freely available for analysis.

## Authors’ contributions

Experimental design: HJC, EBW, GPC

Implementation: HJC, EBW, JN

Clinical support including surgery and sample provision: SCS, AAM, RKM, RC

Analysis and interpretation of the data: HJC, EBW, GPC, LFS, JN, AD, HW, JN-B, SCS, AA

GSC provision: HW

Informatics and statistical analyses: LFS, JN, KAR, AD and GPC

Manuscript preparation: HJC, GPC and EBW, with input from all authors.

