## Supplementary Figures for "Expression profiling of single cells and patient cohorts identifies multiple immunosuppressive pathways and an altered NK cell phenotype in glioblastoma"

**Supplementary Figure 1**

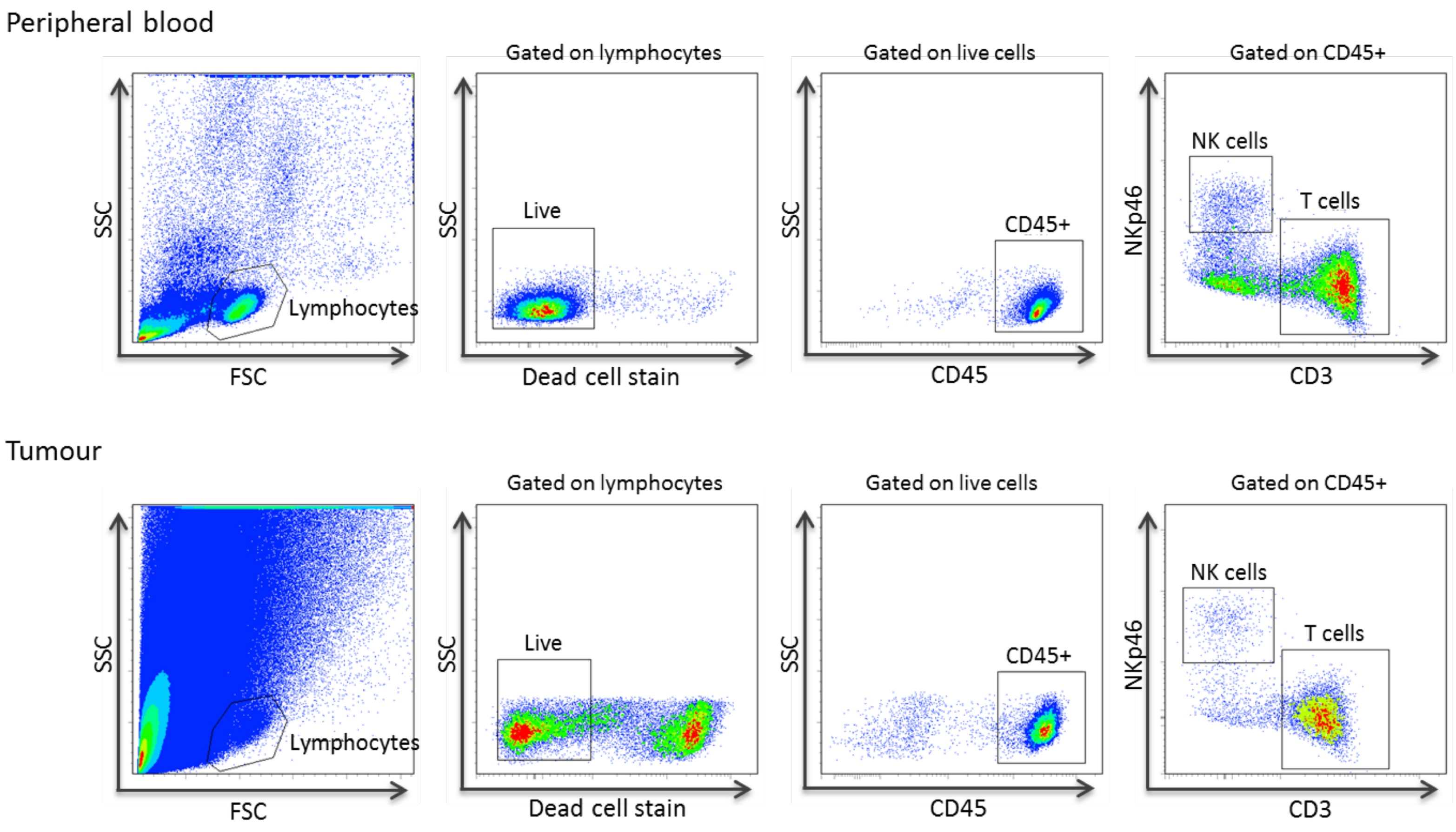


**Gating strategy for identification of T cells and NK cells in blood (top) and tumour (bottom) samples.**
For blood, mononuclear cells were isolated on a Ficoll gradient and for tumour samples, single cell suspensions were generated using mechanical disruption. Cells were stained with appropriate antibodies and lymphocytes identified using forward and side scatter (FSC, SSC). Live cells in the lymphocyte gate were identified using a dead cell discriminator. Haematopoietic cells were identified via staining with anti-CD45 antibodies and CD45+ cells were then analysed for NK cells (NKp46+CD3negative) and T cells (NKp46negativeCD3+). Note that NKp46 was used to identify NK cells in place of CD56 as many cells in the brain express the CD56 molecule.
These NK cell and T cell gates were used to determine the expression of PD-1, NKG2D etc as shown in the main text.

**Supplementary Figure** 2

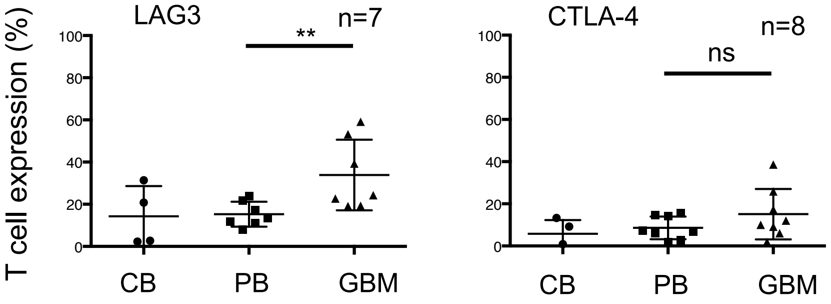


**The cell surface phenotype of GBM-infiltrating lymphocytes**

Expression of LAG3 and CTLA-4 on CD3+ T cells in GBM patient tumour (GBM), patient blood (PB) and control blood from healthy donors (CB). Each dot represents a single patient sample (with number of patients samples n analysed for each sample shown); the bar indicates the mean± standard deviation. The patient-derived tumour (GBM) and blood (PB) samples were analysed using a paired t test; *P<0.05, **P<0.01; ***P<0.001; ns: not significant.

**Supplementary Figure 3**

A

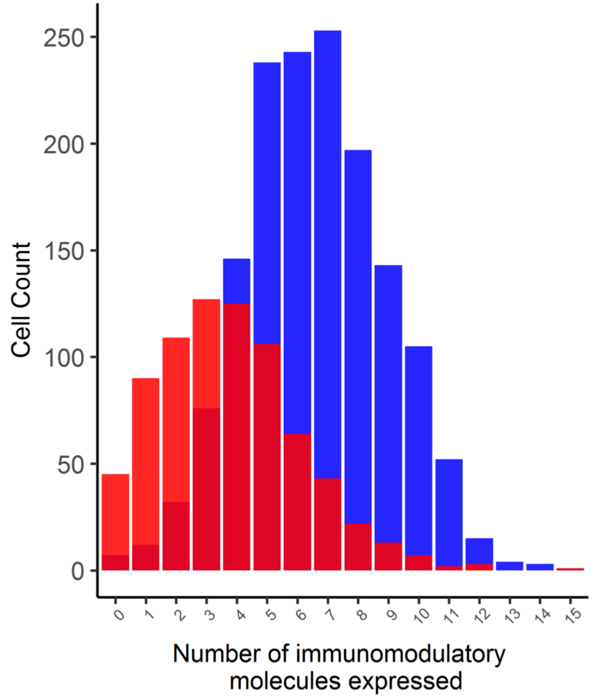


B

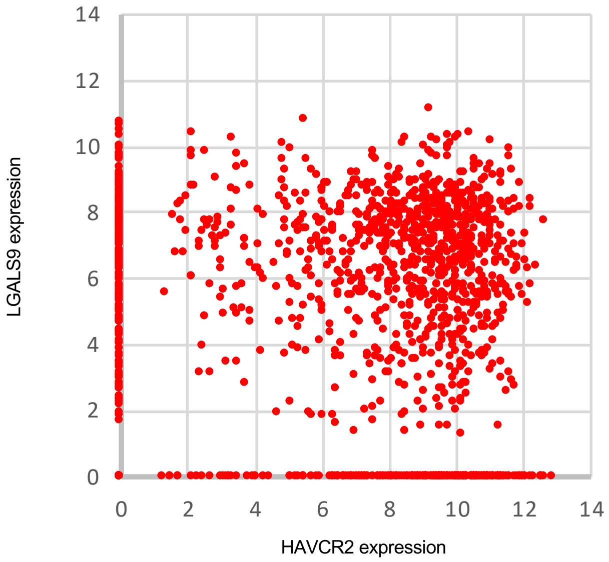


**Expression of immunomodulatory molecules in GBM single cell RNAseq data**

**A)** The number of different candidate immunosuppressive genes expressed by tumour cells (defined as SOX9+EGFR+; shown in red)) and immune cells (defined as PTPRC+; shown in blue).

**B)** Expression of HAVCR2 (encoding the immune checkpoint molecule TIM-3) and LGALS9 (encoding the TIM-3 ligand, galectin-9) amongst PTPRC expressing cells.

### Supplementary Figure 4

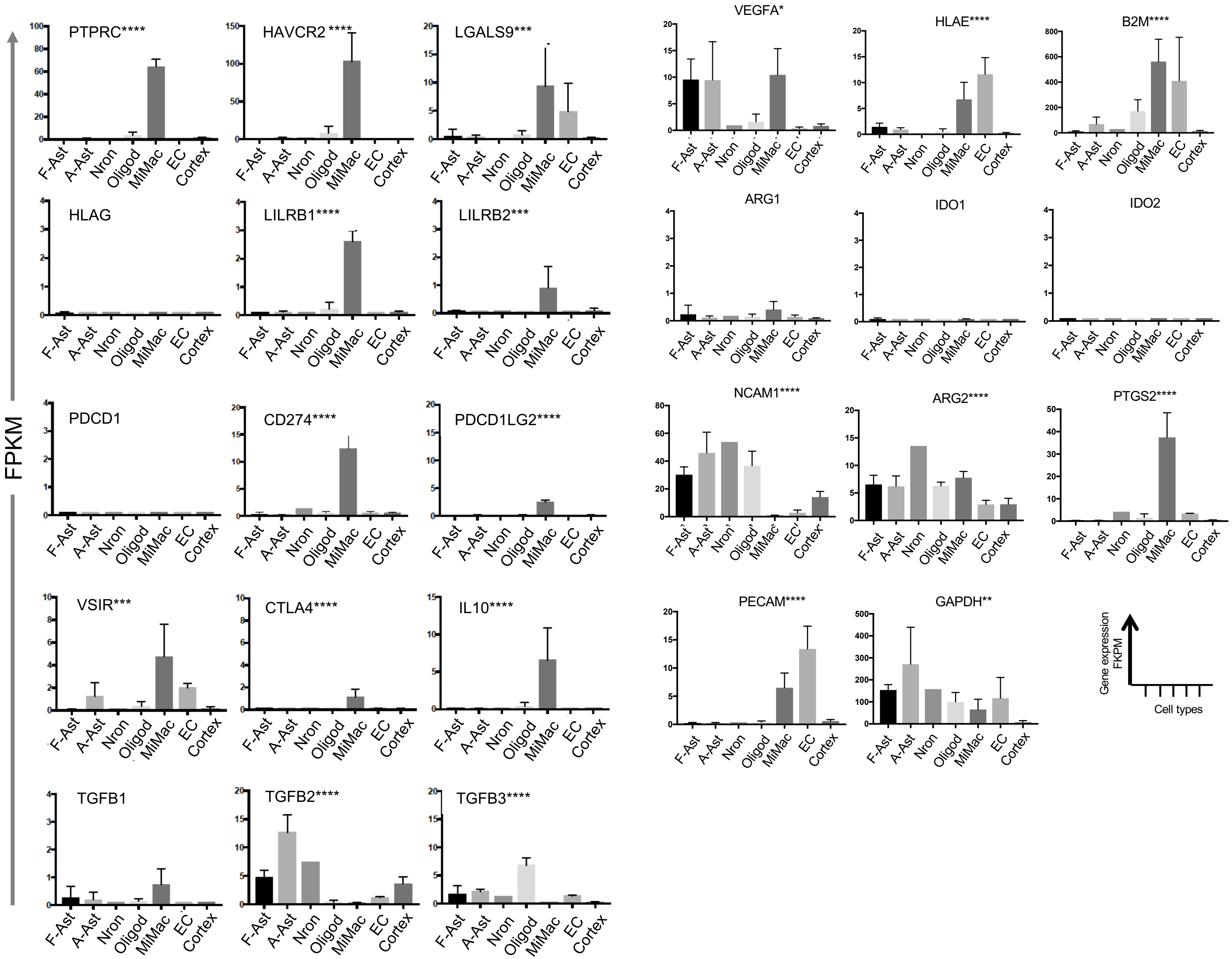


**Gene expression in brain cell populations in the absence of tumour cells.**

Graphs show expression of selected genes expressed as Fragments Per Kilobase of transcript per Million mapped reads (FPKM) across six different brain-derived cell types, with various numbers of donors (n) per cell type; foetal astrocytes (F-Ast, n=6), adult astrocytes (A-Ast, n=12), neurons (Nron, n=1), oligodendrocytes (Oligod, n=5), microglia/macrophage (MiMac, n=3), endothelial cells (EC, n=2) and whole cortex (Cortex, n=4). Bars show standard deviation from mean. Gene names are shown with asterisks where gene expression across the panel is significantly different (using one way ANOVA); *P<0.05, **P<0.01, ****P<0.0001. This data was downloaded from the study of Zhang et al (2014). Expression of any gene can be viewed within this dataset using the tool at brainrnaseq.org.

**Supplementary Figure 5.**

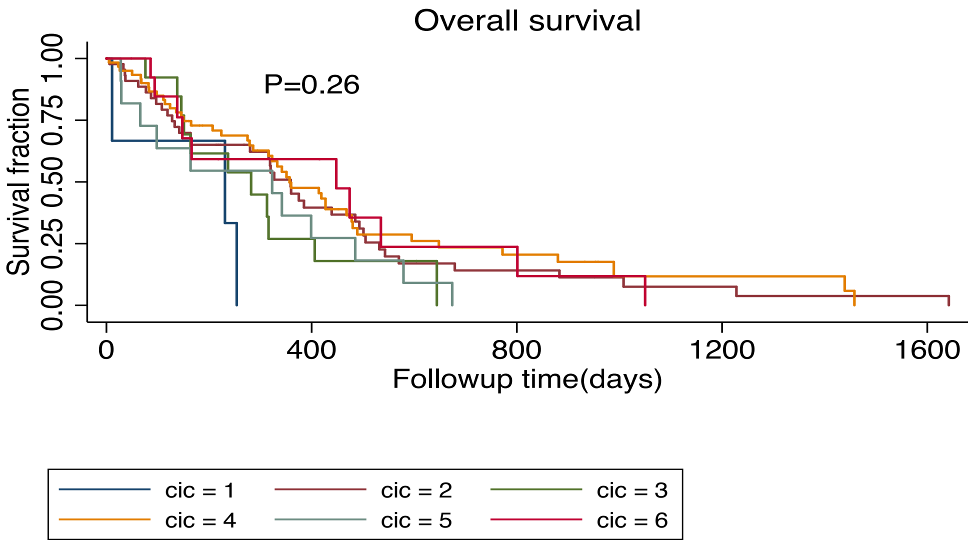


**Survival of GBM patients classified according to CIC**

Kaplan-Meier plot of GBM patient survival according to CIC classification. Patients were classified into CICs according to the data in Figure 6 of the main manuscript. Survival fraction was determined using the data provided in the TCGA database.

### Supplementary Figure 6

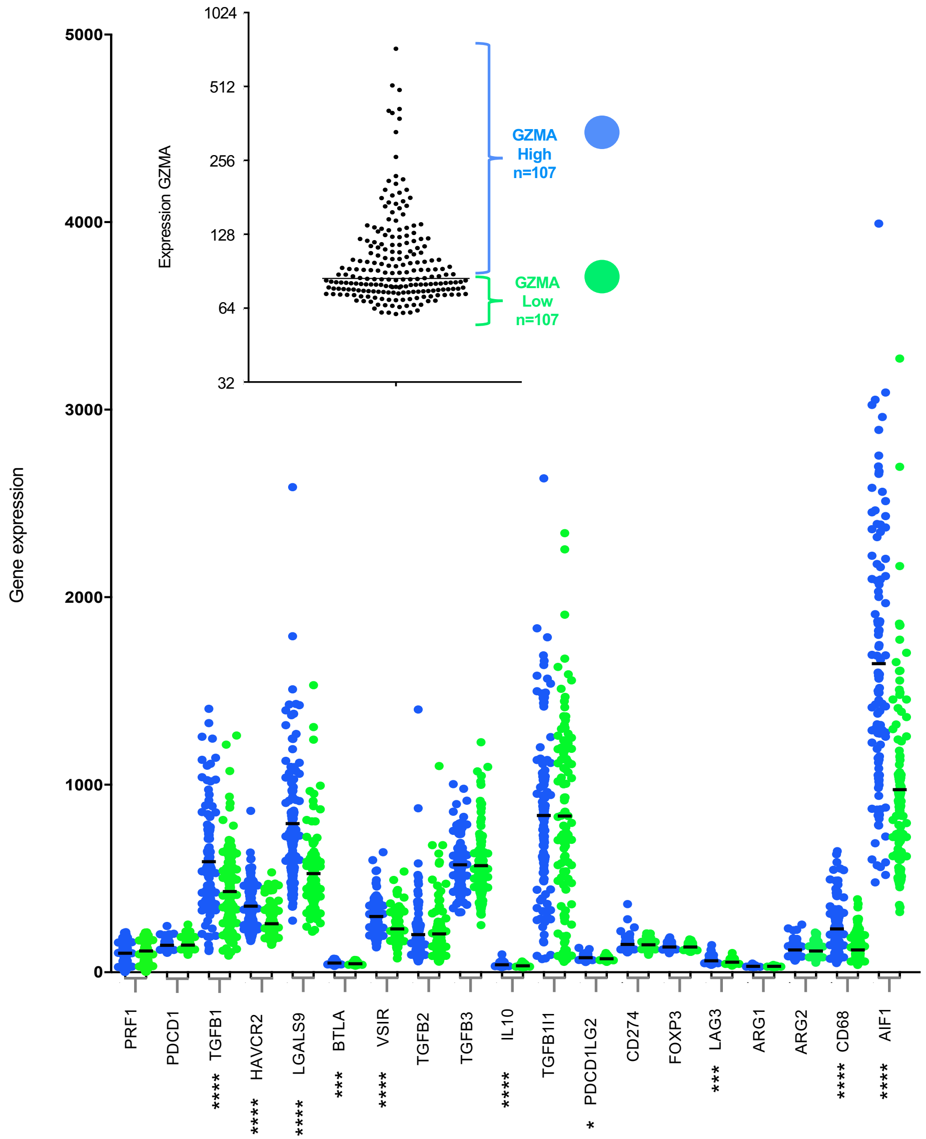


**Microarray data from 214 GBM patients in the REMBRANDT study**

The 214 patients were classified as either high GZMA expressers (n=107) or low GZMA expressers (n=107) by splitting at the median expression value (inset graph). The main graph shows expression of selected genes in the GZMA high (blue) and GZMA low (green) patient groups. Dots represent individual patients, with the mean value indicated by a bar. Each pair of samples was analysed using the Mann Whitney U test. Samples showing statistically different distributions are indicated under the x-axis; *P<0.05, ***P=0.0001, ****P<0.0001. The REMBRANDT gene expression data was downloaded from betastasis.com

| CD specificity | Conjugate | Clone | Manufacturer, product number | Host | Target | Class |
|---|---|---|---|---|---|---|
| CD2 | PE | RPA-2.10 | BD Biosciences, 555327 | Mouse | Human | IgG1k |
| CD3 | PerCP | BW264/56 | Miltenyi Biotec, 130-094-965 | Mouse | Human | IgG2a |
| CD9 | PE | M-L13 | BD Biosciences, 555372 | Mouse | Human | IgG1 |
| CD24 | PE | 32D12 | Miltenyi Biotec, 130-095-953 | Mouse | Human | IgG1 |
| CD45 | FITC | HI30 | BD Biosciences, 555482 | Mouse | Human | IgG1k |
| CD48 | FITC | BJ40 | BioLegend, 336707 | Mouse | Human | IgG1k |
| CD69 | BV421 | FN50 | BioLegend, 310929 | Mouse | Human | IgG1k |
| CD69 | FITC | FN50 | BioLegend, 310904 | Mouse | Human | IgG1k |
| CD90 | VioBlue | DG3 | Miltenyi Biotec, 130-099-271 | Mouse | Human | IgG1 |
| CD107a | PE | H4A3 | BD Biosciences, 555801 | Mouse | Human | IgG1k |
| CD112 (Nectin-2) | PE | R2.525 | BD Biosciences, 551057 | Mouse | Human | IgG1k |
| CD132 (IL-2 Rγ) | PE | AG184 | BD Biosciences, 555900 | Mouse | Human | IgG1k |
| CD152 (CTLA-4) | PE | BNI3 | Miltenyi Biotec, 130-097-684 | Mouse | Human | IgG2a |
| CD184 | PEVio770 | 12G5 | Miltenyi Biotec, 130-103-798 | Mouse | Human | IgG2a |
| CD223 (LAG-3) | PE | REA351 | Miltenyi Biotec, 130-105-452 | Mouse | Human | |
| CD226 (DNAM-1) | PE | DX11 | Miltenyi Biotec, 130-092-476 | Mouse | Human | IgG1 |
| CD244 (2B4) | PE | REA112 | Miltenyi Biotec, 130-099-051 | Mouse | Human | |
| CD273 (PD-L2) | PE | MIH18 | BD Biosciences, 558066 | Mouse | Human | IgG1k |
| CD274 (PD-L1) | PE | MIH1 | BD Biosciences, 557924 | Mouse | Human | IgG1k |
| CD279 (PD-1) | PE | PD1.3.1.3 | Miltenyi Biotec, 130-096-164 | Mouse | Human | IgG2b |
| CD314 (NKG2D) | PE | 1D11 | BD Biosciences, 557940 | Mouse | Human | IgG1k |
| CD317 (Tetherin) | PE | REA202 | Miltenyi Biotec, 130-101-707 | Mouse | Human | |
| CD335 (NKp46) | APC | 9E2/NKp46 | BD Biosciences, 558051 | Mouse | Human | IgG1k |
| CD336 (NKp44) | PE | 2.29 | Miltenyi Biotec, 130-092-480 | Mouse | Human | IgG1 |
| CD337 (NKp30) | PE | P30-15 | BD Biosciences, 558407 | Mouse | Human | IgG1k |
| B7H6 | APC | 875001 | R&D Systems, FAB 7144 | Mouse | Human | IgG1 |
| HLA-A,B,C | APC | W6/32 | BioLegend, 311410 | Mouse | Human | IgG2a |
| MIC A/B | PE | 6D4 | BD Biosciences, 558352 | Mouse | Human | IgG2ak |
| ULBP1 | PE | 170818 | R&D Systems, FAB 1380P | Mouse | Human | IgG2a |
| Isotype | APC | MOPC-21 | BD Biosciences, 555751 | Mouse | | IgG1k |
| Isotype | BV421 | MOPC-21 | BioLegend, 400157 | Mouse | | IgG1k |
| Isotype | FITC | MOPC-21 | BD Biosciences, 555748 | Mouse | | IgG1k |
| Isotype | PE | MOPC-21 | BD Biosciences, 559320 | Mouse | | IgG1k |
| Isotype | PE | G155-178 | BD Biosciences, 555574 | Mouse | | IgG2ak |
| Isotype | PerCP Cy5.5 | MOPC-21 | BD Biosciences, 552834 | Mouse | | IgG1k |
| Isotype | PEVio770 | S43.10 | Miltenyi Biotec, 130-096-638 | Mouse | | IgG2a |
| Isotype | APC | IS5-21F5 | Miltenyi Biotec, 130-092-214 | Mouse | | IgG1k |
| Isotype | VioBlue | IS5-21F5 | Miltenyi Biotec, 130-094-670 | Mouse | | IgG1 |
| Zombie | NIR | | Biolegend, 423106 | | | |

**Supplementary Materials Table** Details of reagents for Flow cytometric analysis.

Supplementary Table 1

GBM4

| CD | %Gated |
|---|---|
| 9 | 99.96 |
| 15 | 28.24 |
| 24 | 99.67 |
| 26 | 45.72 |
| 29 | 94.81 |
| 34 | 54.75 |
| 44 | 97.35 |
| 46 | 99.98 |
| 47 | 100 |
| 54 | 73.14 |
| 55 | 73.42 |
| 56 | 69.62 |
| 57 | 98.98 |
| 59 | 99.98 |
| 61 | 31.87 |
| 63 | 99.92 |
| 71 | 99.75 |
| 73 | 60.17 |
| 81 | 99.8 |
| 90 | 64.35 |
| 91 | 90.62 |
| 94 | 92.85 |
| 95 | 97.53 |
| 97 | 99.99 |
| 98 | 100 |
| 99 | 99.99 |
| 105 | 98.24 |
| 109 | 20.36 |
| 112 | 79.71 |
| 118 | 36.06 |
| 119 | 98.89 |
| 130 | 91.14 |
| 141 | 86 |
| 142 | 92 |
| 146 | 94.54 |
| 147 | 100 |
| 151 | 99.86 |
| 152 | 27.17 |
| 164 | 98.76 |
| 165 | 99.92 |
| 166 | 98.03 |
| 184 | 61.1 |
| 200 | 95.55 |
| 209 | 26.39 |
| 220 | 50.1 |
| 221 | 88.91 |
| 227 | 55.26 |
| 268 | 46.26 |
| 271 | 93.53 |
| 274 | 20.2 |
| 338 | 21.76 |
| 340 | 94.98 |
| 107A | 73.59 |
| 107B | 38.39 |
| 120A | 47.52 |
| 140A | 50.48 |
| 140B | 55.76 |
| 172B | 75.75 |
| 49b | 99.99 |
| 49c | 83.81 |
| 49d | 51.39 |
| 49e | 88.34 |
| 49f | 98.52 |
| 51/61 | 55.64 |
| 99R | 92.65 |
| b2m | 99.81 |
| egfr | 31.04 |
| GD2 | 92.33 |
| HLA ABC | 99.97 |
| HLA DQ | 93.52 |
| HLAA2 | 97.18 |
| MICA/B | 40.37 |
| SSEA1 | 31.86 |

GBM11

| CD | %Gated |
|---|---|
| 9 | 100 |
| 10 | 48.99 |
| 26 | 70.04 |
| 29 | 39.52 |
| 34 | 99.26 |
| 38 | 36.44 |
| 40 | 34.05 |
| 44 | 100 |
| 46 | 100 |
| 47 | 99.99 |
| 54 | 99.41 |
| 55 | 98.61 |
| 56 | 99.75 |
| 57 | 88.91 |
| 59 | 100 |
| 61 | 91.58 |
| 63 | 99.96 |
| 71 | 99.99 |
| 73 | 99.98 |
| 74 | 95.29 |
| 77 | 85.22 |
| 80 | 28.97 |
| 81 | 99.84 |
| 90 | 30.33 |
| 91 | 99.32 |
| 94 | 75.71 |
| 95 | 98.11 |
| 97 | 99.98 |
| 98 | 100 |
| 99 | 99.86 |
| 104 | 98.99 |
| 105 | 48.77 |
| 106 | 98.22 |
| 109 | 29.73 |
| 112 | 32.23 |
| 119 | 99.06 |
| 130 | 56.11 |
| 141 | 61.41 |
| 142 | 78.23 |
| 146 | 96.73 |
| 147 | 99.99 |
| 151 | 88.73 |
| 164 | 99.98 |
| 165 | 99.67 |
| 166 | 99.68 |
| 184 | 43.79 |
| 201 | 40.7 |
| 205 | 99.62 |
| 221 | 37.27 |
| 227 | 95.48 |
| 268 | 41.13 |
| 271 | 98.13 |
| 273 | 81.29 |
| 274 | 97.28 |
| 321 | 20.37 |
| 338 | 38.87 |
| 340 | 86.48 |
| 107A | 99.4 |
| 107B | 98.22 |
| 120A | 93.31 |
| 121B | 23.32 |
| 140A | 23.9 |
| 172B | 34.05 |
| 49b | 99.97 |
| 49c | 100 |
| 49d | 96.86 |
| 49e | 100 |
| 49f | 99.67 |
| 51/61 | 96.93 |
| 62P | 20.62 |
| 99R | 88.77 |
| b2m | 99.96 |
| GD2 | 99.94 |
| HLA ABC | 99.97 |
| HLA DQ | 99.46 |
| HLA DR | 99.98 |
| HLA DRPQ | 99.95 |

GBM13

| CD | %Gated |
|---|---|
| 9 | 100 |
| 24 | 78.56 |
| 26 | 70.49 |
| 29 | 91.72 |
| 34 | 55.04 |
| 44 | 100 |
| 46 | 100 |
| 47 | 99.98 |
| 50 | 21.35 |
| 54 | 94.73 |
| 55 | 100 |
| 56 | 99.56 |
| 57 | 96.16 |
| 59 | 99.95 |
| 61 | 84.77 |
| 63 | 99.92 |
| 71 | 99.55 |
| 73 | 99.97 |
| 77 | 64.76 |
| 80 | 25.44 |
| 81 | 99.83 |
| 90 | 99.06 |
| 91 | 99.25 |
| 94 | 97.98 |
| 95 | 99.76 |
| 97 | 100 |
| 98 | 100 |
| 99 | 99.99 |
| 104 | 70.65 |
| 105 | 88.5 |
| 108 | 65.92 |
| 112 | 71.2 |
| 118 | 44.6 |
| 119 | 98.85 |
| 130 | 93.38 |
| 141 | 98.98 |
| 142 | 99.19 |
| 144 | 41.24 |
| 146 | 98.83 |
| 147 | 99.99 |
| 151 | 99.89 |
| 152 | 37.74 |
| 164 | 96.08 |
| 165 | 99.98 |
| 166 | 99.47 |
| 184 | 56.94 |
| 200 | 98.67 |
| 209 | 61.79 |
| 220 | 81.34 |
| 221 | 78.72 |
| 227 | 62.9 |
| 268 | 50.41 |
| 271 | 99.87 |
| 274 | 75.63 |
| 321 | 84.49 |
| 329 | 34.7 |
| 338 | 38.32 |
| 340 | 97.36 |
| 107A | 88.26 |
| 107B | 54.28 |
| 120A | 71.32 |
| 140B | 37.51 |
| 172B | 89.85 |
| 49a | 35.63 |
| 49b | 99.98 |
| 49c | 100 |
| 49d | 23.75 |
| 49e | 99.93 |
| 49f | 99.77 |
| 51/61 | 93.03 |
| 99R | 99.45 |
| b2m | 99.94 |
| CD326 | 63.61 |
| egfr | 53.2 |
| GD2 | 99.96 |
| HLA ABC | 99.99 |
| HLA DQ | 99.72 |
| HLAA2 | 99.22 |
| MICA/B | 43.86 |
| SSEA4 | 32.18 |

GBM1

| CD | %Gated |
|---|---|
| 9 | 26.7 |
| 15 | 36.8 |
| 26 | 32 |
| 29 | 62 |
| 34 | 3.8 |
| 44 | 71.6 |
| 46 | 89.3 |
| 47 | 91.6 |
| 54 | 23.6 |
| 56 | 82.7 |
| 57 | 71.3 |
| 58 | 77.2 |
| 58 | 77.2 |
| 59 | 80.9 |
| 63 | 83.2 |
| 71 | 87.1 |
| 73 | 37.9 |
| 81 | 89.8 |
| 90 | 81.3 |
| 91 | 57.7 |
| 94 | 70.5 |
| 95 | 73.8 |
| 97 | 86 |
| 98 | 75.6 |
| 99 | 87.2 |
| 106 | 25.6 |
| 119 | 75.1 |
| 142 | 82.6 |
| 146 | 83.3 |
| 147 | 90.9 |
| 151 | 89 |
| 164 | 76.7 |
| 165 | 87.8 |
| 166 | 87.3 |
| 200 | 70.6 |
| 227 | 50.5 |
| 271 | 53.4 |
| 321 | 21.5 |
| 340 | 26.8 |
| 107a | 41.4 |
| 140a | 28.2 |
| 49a | 34.9 |
| 49b | 89.1 |
| 49c | 29.2 |
| 49d | 31 |
| 49e | 68.6 |
| 49f | 63.8 |
| 99r | 36.7 |
| b2m | 83.8 |
| gd2 | 53.8 |
| HLA ABC | 90.8 |
| HLA DQ | 40.3 |
| ssea1 | 58.9 |

GBM20

| CD | %Gated |
|---|---|
| 9 | 99.97 |
| 15 | 67.12 |
| 24 | 89.22 |
| 26 | 55.02 |
| 29 | 65.05 |
| 34 | 56.14 |
| 44 | 99.84 |
| 46 | 99.98 |
| 47 | 100 |
| 50 | 31.08 |
| 54 | 38.51 |
| 55 | 34.17 |
| 56 | 99.75 |
| 57 | 99.75 |
| 58 | 100 |
| 59 | 99.92 |
| 61 | 88.07 |
| 63 | 99.95 |
| 71 | 99.64 |
| 73 | 99.87 |
| 74 | 37.5 |
| 75 | 22.84 |
| 77 | 65.45 |
| 80 | 57.11 |
| 81 | 100 |
| 88 | 47.35 |
| 90 | 99.04 |
| 91 | 94.09 |
| 94 | 89.63 |
| 95 | 96.69 |
| 97 | 99.97 |
| 98 | 100 |
| 99 | 99.99 |
| 105 | 74.88 |
| 106 | 45.03 |
| 108 | 75.15 |
| 112 | 55.1 |
| 118 | 20.41 |
| 119 | 98.14 |
| 130 | 98.03 |
| 138 | 47.21 |
| 141 | 94.31 |
| 142 | 97.51 |
| 146 | 96.92 |
| 147 | 100 |
| 151 | 99.98 |
| 152 | 75.76 |
| 164 | 97.38 |
| 165 | 99.97 |
| 166 | 94.15 |
| 171 | 36.09 |
| 184 | 47.56 |
| 200 | 98.92 |
| 205 | 20.88 |
| 209 | 47.54 |
| 220 | 81.45 |
| 221 | 58 |
| 227 | 91.57 |
| 231 | 21.87 |
| 268 | 73.6 |
| 271 | 98.5 |
| 273 | 34.4 |
| 274 | 72.22 |
| 321 | 42.19 |
| 338 | 90.98 |
| 340 | 79.74 |
| 107A | 78.66 |
| 107B | 71.14 |
| 120A | 69.01 |
| 120B | 41.99 |
| 140A | 26.31 |
| 140B | 66.96 |
| 172B | 70.02 |
| 49b | 99.33 |
| 49c | 99.96 |
| 49d | 82.31 |
| 49e | 99.83 |
| 49f | 93.66 |
| 51/61 | 94.17 |
| 99R | 97.86 |
| b2m | 100 |
| egfr | 68.68 |
| GD2 | 99.82 |
| HLA ABC | 99.95 |
| HLA DQ | 98.28 |
| HLA DR | 96.45 |
| HLA DRPQ | 95.51 |
| HLAA2 | 99.94 |
| MICA/B | 77.53 |
| SSEA1 | 66.06 |

Full list of antigen screened can be found
http://www.bdbiosciences.com/ds/pm/others/23-10930.pdf
Data shows percentage expressed of each antigen (CD).

**Supplementary Table 2**

| Immune Group | Classical | Mesenchymal | Proneural | *Total* |
|---|---|---|---|---|
| **CIC1** | 0 | 1 | 2 | *3* |
| **CIC2** | 13 | 24 | 7 | *44* |
| **CIC3** | 4 | 7 | 1 | *12* |
| **CIC4** | 29 | 5 | 29 | *63* |
| **CIC5** | 3 | 6 | 2 | *11* |
| **CIC6** | 6 | 6 | 2 | *14* |
| *Total* | *55* | *49* | *43* | *147* |

**The number of TCGA patient tumours classified into each GBM expression subtype per immune group.** A chi-squared test on the distributions within the largest immune groups, CIC2 and CIC4, show that these are significantly different ($p=3.8\times10^{-7}$), with the mesenchymal subtype enriched in the former and depleted in the latter
